# Quantifying Spatio-Temporal Overlap of Invasive Wild Pigs and Domestic Pig Farms as a Proxy for Potential Disease Transmission Risk

**DOI:** 10.1101/2022.09.25.509397

**Authors:** Ruth A. Aschim, Ryan K. Brook

## Abstract

Direct and indirect interactions between livestock and free-ranging wildlife creates important risks to animal health and agricultural productivity. The interface between newly established and rapidly spreading invasive wild pigs and the 2,549 domestic pig farms on the Prairie Provinces of western Canada has created important but poorly understood disease transmission risks. We mapped the spatial overlap of wild and domestic pigs to identify the areas of highest risk and associated distribution of diseases of concern using databases of wild pig occurrences and domestic pig farm locations. We also examined spatial and temporal overlap at the individual farm scale using GPS collared invasive wild pigs. Across the provinces of Alberta, Saskatchewan, and Manitoba, spatial overlap of invasive wild pigs with all combined, large-scale domestic pig farms, small-scale domestic pig farms, and domestic wild boar farms was 21%, 21%, 21%, and 53%. Invasive wild pig locations were significantly closer to domestic pig farms and domestic wild boar farms compared to random points on the landscape. The number of wild pig occurrences was greatest within 20 km of domestic pig farms and decreased linearly as distance increased. The Canadian distribution of wild pigs had considerable spatial overlap with recent areas detected with bovine tuberculosis (6,002 km^2^) in livestock and wildlife and Chronic Wasting Disease (156,159 km^2^) in wildlife, including mule deer, white-tailed deer, elk, and moose. The single best predictor of invasive wild pig occurrences across the landscape was close proximity to current or recently past existing domestic wild boar farms. The distance of GPS-collared wild pigs was significant for sex, farm type, month, and season and in southeastern Saskatchewan, average distance to domestic pig farms was 5.3 km. The weighted sum of cover type proportions, wild pig distance to domestic pig and wild boar farms, farm type, and farm density identified the relative risk of wild pig presence associated to each domestic pig farm occupied watershed. Risk was highest for small-scale domestic pig farms and lowest for large-scale domestic pig farms. Our findings highlight important potential routes for disease transmission at the invasive wild pig-domestic pig interface and identify areas where biosecurity improvements are urgently needed. While complete eradication of invasive wild pigs in Canada is no longer achievable, improved passive and active monitoring and removal of wild pigs is critical, especially where risks to domestic pig herds is highest.

## 1. Introduction

Anthropogenic land-use changes have caused significant wildlife habitat loss and fragmentation globally (Hassell et al., 2017). Encroachment of anthropogenic land-use onto wildlife habitat has led to an increase in human-wildlife conflicts in agriculture dominated landscapes at the wildlife-livestock interface (Brook, 2009; Brook and McLachlan, 2009; Rhyan and Spraker, 2010; Simbieda et al., 2011; Brook et al., 2013). The increasing frequency and duration of interactions at the wildlife-livestock interface and the creation of new junctures has created additional avenues of contact between wildlife and livestock, facilitating direct and indirect disease transmission risk (Rhyan and Spraker, 2010; Simbieda et al., 2011).

The incidence of emerging zoonotic and animal diseases has increased through the 21^st^ century and directly corresponds with the increased wildlife component at the wildlife-livestock interface (Jones et al., 2008). Wildlife play a central role in the introduction, maintenance, and transmission of reportable domestic animal diseases (Miller et al., 2017). Indeed, in North America, 79% of reportable animal diseases have identified a wildlife component; 38% of which affect multiple species of livestock and 40% of which are zoonotic (Miller et al., 2013). Interactions at the livestock-wildlife interface pose a continued threat for disease introduction, maintenance, and spread as the livestock-wildlife interface is dynamic and multi-faceted with opportunities for bi-directional pathogen transmission between wildlife and livestock (Bengis et al., 2002; Wiethoelter et al., 2015). On-farm management practices can contribute to livestock-wildlife interactions from access to stored high-quality feed, spilled feed, mineral supplements, and attractive odors (Wycoff et al., 2009; Brook et al., 2013; Pruvot et al., 2014). Complex management challenges, health and safety concerns, and threats to economic sustainability arise with the presence of wildlife hosts and reservoirs of infectious disease (Profitt et al., 2011; Miller et al., 2013).

A barrier to disease eradication in livestock exists with wildlife hosts and reservoirs on the landscape due to the continued and increased interactions at the wildlife-livestock interface (Witmer et al., 2003; Miller et al., 2013). A lack of baseline knowledge on the presence and prevalence of disease in wildlife populations, difficulty locating infected individuals, unknown or poor wildlife population estimates, and the high costs associated with implementing effective vaccines or depopulation efforts compounds the challenges associated with complete disease eradication (Corner, 2006; Olsen, 2010; Rhyan and Spracker, 2010; Miller and Sweeny, 2013). Additionally, the epidemiology of wildlife disease is such that the environmental lifespans of pathogens, the susceptibility of individuals, and routes of transmission differ greatly which makes risk analysis, surveillance, and mitigation challenging (Ryan and Spracker, 2010; World Organization for Animal Health, 2010). These challenges associated with eliminating disease in wildlife populations pose a continued threat of disease (re)-introduction to livestock and has the potential for spill-over back into wildlife populations (Corner, 2006; Olsen, 2010; Miller and Sweeny, 2013).

The spatial distribution, movement, and social processes of wildlife populations are central to disease system dynamics as they can determine how far the disease can spread and influence direct and indirect transmission events (Robinson et al., 2013; Podgorski et al., 2018; Pepin et al., unpublished). Wildlife movement patterns vary temporally (annually, seasonally, and daily) and the factors influencing movement patterns are dependent on a multitude of components, such as the distribution of resources across the landscape, habitat fragmentation, population density, and farm management practices (Manley et al., 1993; Guisan and Thullier, 2005; Brook et al. 2012; Guisan et al., 2017). Additionally, routes of pathogen transmission at the wildlife-livestock interface are often poorly understood (Witmer et al., 2003; Miller et al., 2013) with interactions between wildlife and livestock occurring via direct contact through farm visitations (Wyckoff et al., 2009) and along fence lines (Bengis et al., 2002) or by indirect contact through shared resource use (Siembieda et al., 2011), contamination of water and feed (Sorensen et al., 2014), and infected carcasses (Probst et al., 2017; Fischer et al., 2020). As such, the ability to accurately quantify wildlife-livestock interactions at large scales is limited, and often relies on proxy measures to estimate potential disease transmission risk such as spatial overlap and contact rates (Brook and McLachlan, 2009; Brook et al. 2013). Identifying the resource use of wildlife at the landscape level is fundamental to determining spatial patterns across the landscape. Species-environment relationships determine spatial patterns and movements of species within their geographical range and how they use resources and occupy ecological niches within their home ranges (Fletcher and Fortin, 2019). These relationships remain less clear for invasive species.

Although a relatively new invasive species on the Canadian landscape, invasive wild pigs are well-established and widespread across a large and rapidly expanding area of Canada (Aschim and Brook, 2019). Invasive wild pigs in Canada are hybrids between European wild boar (*Sus scrofa*) and domestic swine (Sus *scrofa domesticus*) (Government of Saskatchewan, 2001; Brook, unpublished). The species is susceptible to 35% of reportable diseases defined by the Canadian Food Inspection Agency (CFIA) (2020). Diseases of concern at the livestock-wildlife interface between wild and domestic pigs include swine brucellosis (*Brucellosis suis*), bovine tuberculosis (*Mycobacterium bovis*), pseudorabies (Aujeszky’s disease) (*Herpesevirius spp*.), Porcine Reproductive and Respiratory Syndrome Virus (PRRSV), and African Swine Fever (ASF) (West et al., 2009; Barrios-Garcia and Ballari, 2012, Bosch et al., 2016; Guinat et al., 2016; Canadian Food Inspection Agency, 2020). In many areas of their native and introduced range wild pigs are disease reservoirs and maintenance hosts due to their high densities, complex social behaviours, and ability to maintain pathogen(s) without a continued source of infection (Naranjo et al., 2008; Ruiz-Fons et al., 2008; Olsen, 2010). Therefore, wild pigs play a significant epidemiological role in the transmission and spread of pathogens across their native introduced range.

The presence of invasive wild pigs on the landscape poses new and increased risks of disease transmission and spread to the domestic swine industry across Canada. Disease threats are a significant concern to livestock producers and industries as disease outbreaks are associated with high economic losses from livestock morbidity and mortality, trade implications, and the increased costs associated treatment and controls (Witmer et al., 2003; Seward et al., 2004; Siembieda et al., 2011; Barrios-Garcia and Ballari, 2012; Brown et al., 2020). Domestic swine in Canada are currently free from all reportable diseases (Canadian Food Inspection Agency, 2020), which allows for international trade and is what has established Canada as a leading pork exporter globally (Agri and Agri-Food Canada, 2015; Statistics Canada, 2015). The recently established invasive wild pig population, however, poses new threats and challenges to this disease-free status. The pork industry is a significant contributor to the Canadian economy as the fourth largest agricultural commodity in Canada, accounting for 30% of all Canadian livestock exports and over $4.2 billion annually in exports (Agri and Agri-Food Canada, 2021; Canadian Pork Council, 2021; Canada Pork International, 2021). Thus, the presence of wild pigs in Canada as potential disease vectors poses both economic and epidemiologically significant risks to the pork industry and the Canadian economy.

The presence of invasive wild pigs on the landscape compounds the existing challenges associated with wildlife disease management and poses new and continued risks of pathogen introduction, transmission, and spread between wild and domestic pigs. As such, risk analyses and mitigation strategies that prevent wildlife-livestock spatial overlap are fundamental to the prevention of disease transmission at the wildlife-livestock interface (Miller et al., 2013). A central challenge in mitigating the risks posed to domestic pigs in Canada, however, is that the current and likely future spatial-temporal overlap of wild pigs with domestic pig farms is unknown. Due to the complexities of disease transmission dynamics, the widespread invasive wild pig population, and the broad geographic distribution of domestic pig farms, the ability to accurately identify and quantify direct and indirect contact between wild and domestic pigs is a logistical challenge (Brook and McLachlan, 2009). As such, understanding the spatial patterns and resource use of invasive wild pigs based on landscape composition and livestock demographics can provide insight into areas where spatial and temporal overlap between wild and domestic pigs may occur, increasing the potential for disease transmission and spread (Brook and McLachlan, 2009; Berentsen et al., 2014; Carrasco-Garcia et al., 2016). The objectives of this study were to: (1) identify areas and times where wild and domestic pig spatial overlap occurs at watershed and individual farm scales, (2) identify environmental variables that best predict where spatial overlap occurs at multiple spatial scales, (3) map the distribution of identified key diseases of concern that occur within the range of wild pigs in Canada and the United States, and (4) use spatial overlap as a proxy for potential disease transmission risk to support what hopefully one day would be a systematic disease monitoring and mitigating strategy in Canada.

## 2. Materials and Methods

### 2.1 Study Area

The study area included the three Canadian Prairie Provinces of Alberta, Saskatchewan, and Manitoba where domestic and wild pigs have been detected. The overall area is within an agriculture dominated landscape and was delineated by the distribution of domestic pig farm occupied watersheds at the Level 9 watershed unit (Verdin and Verdin, 1999; Aschim and Brook 2019). The distribution of domestic pig farm occupied watersheds encompassed an area of 331,543 km^2^ and included the Prairie, Boreal Plains, Boreal Shield, and Montane Cordillera ecozones (49°N-59°N; 120°W-95°W). More than two thirds of the study area (69%) fell within the Prairie Ecozone. The continental inland location of the Prairie Ecozone was characterized by an extreme climate, with long cold winters and short hot summers (Shepard and McGinn, 2003). Mean annual temperatures in the coldest and warmest months ranged from −7.8°C to −17.8 °C, and 15.5 °C and 19.5 °C in Lethbridge and Winnipeg respectively (Natural Resources Canada, 2015). Precipitation levels were low in the semiarid grasslands of the Prairie ecozone with annual precipitation levels below 300mm (Natural Resources Canada, 2015). Precipitation increased both west towards the Rocky Mountains and north to the boreal forest with annual precipitation levels over 1000 mm and 700 mm respectively (Natural Resources Canada, 2015). However, these patterns have been changing in response to climate change.

The western Canadian prairies are the most extensively altered ecozone in Canada (Natural Resources Canada, 2015) and accounted for 80% of Canada’s agricultural land base (Sauchyn and Kulshreshth, 2008). Landscape composition within the study area consisted primarily of agricultural land use (65%), grassland (14%), water (8%), forest cover (12%), and anthropogenic development (1%). Domestic pig production within the prairies constituted a large portion of the provincial agricultural sectors. Manitoba, Alberta, and Saskatchewan are the third, fourth, and fifth largest swine producers in Canada, with a total of 3,345,000, 1,565,000, and 935,000 swine on farms respectively (Saskatchewan Pork Board, 2021; Statistics Canada, 2021). The study area that was used to develop the resource selection function (RSF) model was delineated by wild pig occupied watersheds at the Level 9 watershed unit (Verdin and Verdin, 1999). The output of the RSF model was extrapolated to the extent of all domestic pig farm occupied watersheds in the Prairie Provinces to visualize the potential risk to all current domestic swine operations within Alberta, Saskatchewan, and Manitoba (Fig. 1). As a result, some domestic pig farm occupied watersheds are slightly outside of the convex hull study area of wild pig occurrences used to develop the RSF.

**Figure 1.**
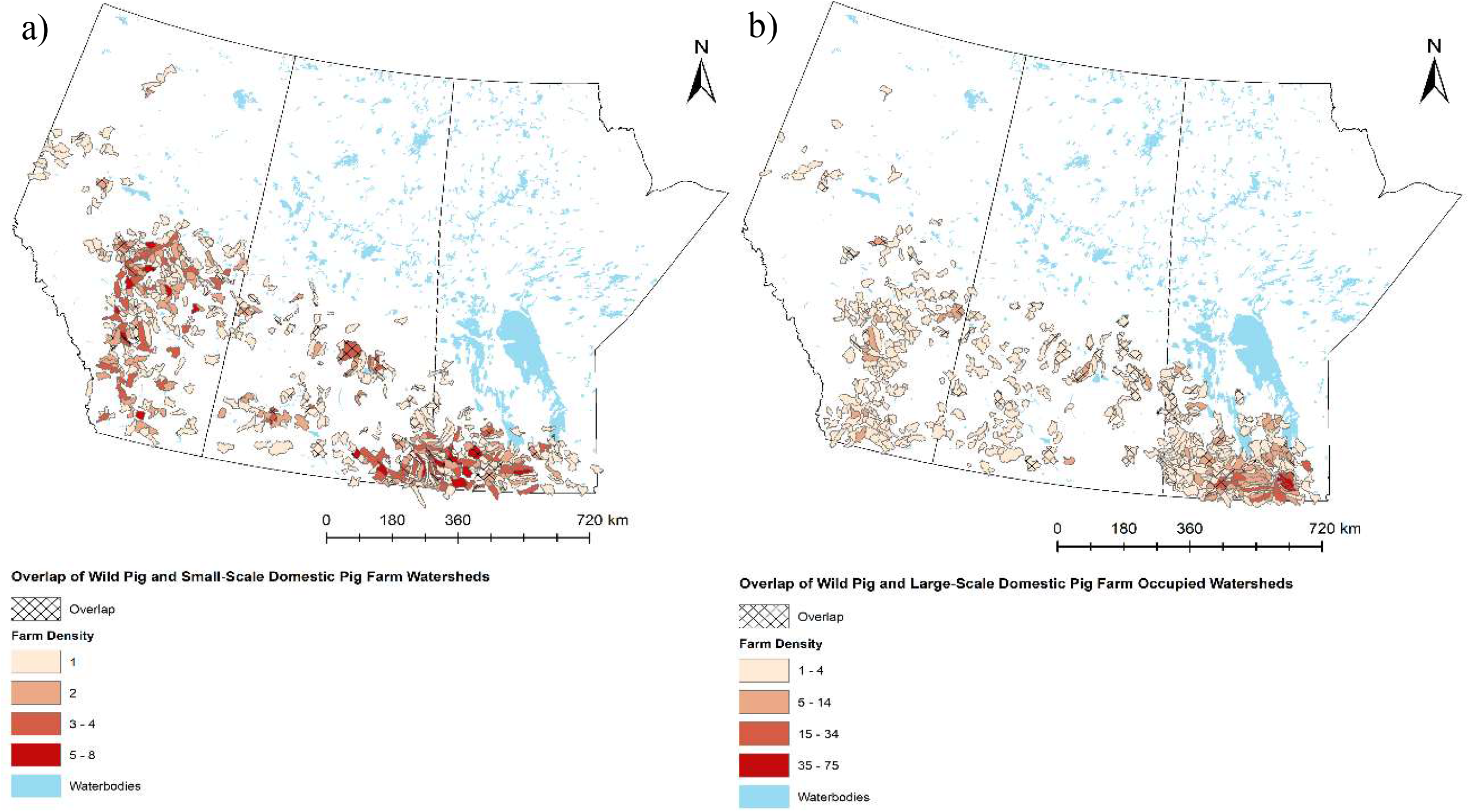
The study area for the resource selection function (RSF) established from a convex hull of all recorded wild pig locations (n=873) at the Level 9 watershed unit across Alberta, Saskatchewan, and Manitoba from 1990-2017.

Domestic swine farm densities differ across the three provinces, with Saskatchewan having a relatively lower concentration of swine operations per square km of arable land at 42.2 as compared to Alberta and Manitoba which have densities of 100 and 188.7 domestic swine operations per square km respectively. Across their Canadian range, 99% of all wild pig occurrences were located within the Prairie Provinces (Aschim and Brook, 2019), therefore, the overwhelming majority of the risk of spatial overlap of invasive wild pigs with domestic pig farms is concentrated within this area.

### 2.2 *Invasive Wild Pig and Domestic Pig Farm Distribution* Mapping

Canadian wild pig locations were identified through surveys with wildlife professionals and stakeholders, media reports, and trail camera data (Aschim and Brook, 2019). Locations were obtained as a Legal Land Location, UTM coordinate, or distance and direction from nearest town. All locations were converted to UTM locations using Google Maps (2018) and the Legal Land Description Converter (2017). Point UTM locations were aggregated up to the Level 9 watershed unit (x area = 371 km^2^) following the Pfafstetter classification system (Verdin and Verdin, 1999). The social survey sampling design used to collect wild pig occurrence data was approved by and conducted in accordance with local ethical review [SPECIFIC DETAILS REMOVED FOR DOUBLE BLIND PEER REVIEW].

Domestic pig farm location data was obtained from the Alberta, Saskatchewan, and Manitoba Pork Boards. Locations were provided in the form of legal land descriptions (LLD) and the farm was described as either a large or small-scale operation. Information on the number of animals that classified a large versus a small-scale operation was not provided. Large-scale operations were classified as biosecure facilities housing large numbers of pigs at various stages of farrowing to finishing, whereas small-scale operations were considered backyard or hobby farms with small numbers of pigs and no biosecurity measures. Due to confidentiality concerns, domestic pig farm locations were aggregated up to the Level 9 watershed scale. Occupied watersheds were classified into three different datasets: all domestic pig farms combined, large-scale domestic pig farms, and small-scale domestic pig farms. The distinction was made between these farm types to determine if the risks posed by wild pigs differed between classifications. Separate maps were created that examined the spatial overlap of domestic pig farms and wild pig occupied watersheds for large and small-scale domestic pig farms. The density of domestic pig farms within wild pig occupied watersheds was identified and the area of spatial overlap between wild pigs and domestic pig and wild boar farms was quantified. The mean, maximum, and minimum distances between wild pig UTM locations and the UTM centroid from domestic pig and domestic wild boar farm LLD’s were calculated.

The distance (km) of random and used wild pig locations at a 5:1 ratio was examined via boxplots and Welch’s two-sided t-tests to determine if wild pig distance to domestic pig farms (large and small) and domestic wild boar farms (current and historic) was significant compared to random points on the landscape. Used wild pig locations were evaluated using boxplots and Welch’s t-tests to determine if there was a significant difference in the distance (km) of wild pigs to large versus small domestic pig farms. The distance of used wild pig locations to domestic pig versus wild boar farms were also evaluated using the same methods. The relationship between the number of wild pig locations and the distance (km) to domestic pig farms was evaluated using a non-linear regression.

### 2.3 Maps of Pathogens Established in Wildlife Populations across Canada and the U.S

Pathogens established in wildlife populations that are; i) transmissible to wild pigs, or wild pigs are a known pathogen reservoir/host; ii) have serious economic impacts on domestic livestock production (i.e., CFIA reportable diseases); and iii) are known to be established in wildlife populations were identified and mapped across Canada and the U.S. A review of published literature and federal, provincial, and state government websites provided spatial data for areas where selected pathogens are established in wildlife populations. Areas where specific pathogens have been identified in wildlife populations were mapped via county, park, wildlife management unit, and state boundaries, dependent on the available data from each resource. The maps of wild pig distribution and pathogens established in wildlife populations were overlaid in ArcGIS 10.6 (ESRI Redlands CA., 2018) to identify the spatial aspects of the risk of disease transmission to wild pigs from wildlife.

The Canadian map of wild pig distribution from Aschim and Brook (2019) was expanded to include U.S. locations to examine the spatial aspects of wild pig distribution and areas with selected pathogens established in wildlife populations. Canadian locations were identified through systematic and snowball sampling of wildlife professionals and stakeholders (Aschim and Brook 2019), buffered by 10km, and aggregated up to the Level 9 watershed unit following the Pfstatter classification system. U.S. wild pig locations were obtained through federal and state natural resource or equivalent departments in which wildlife management is within the scope of the corresponding jurisdiction. U.S. wild pig locations were mapped at the county level as this was the smallest scale of data that was uniformly available across all states. U.S. locations were not converted into watershed units as the average county size across the U.S. is 1,610 km^2^ (U.S. Census, 2000), which is a significantly larger area than the Level 9 watershed scale used in the Canadian distribution map. Selecting watersheds that intersected counties with wild pig presence to create a contiguous map with Canada would have resulted in over-sampling of the U.S. wild pig distribution.

### 2.4 Habitat and Resource Selection by Invasive Wild Pigs

We employed a use-availability design (Manley et al., 1993) to quantify wild pig resource selection across a set of biotic and abiotic covariates at the third order of selection (Johnson, 1980). A resource selection function (RSF) model was conducted at the scale of Level 9 watershed units across the three Prairie Provinces. The study area for the RSF was established from a convex hull of all recorded wild pig locations at the Level 9 watershed unit across Alberta, Saskatchewan, and Manitoba from 1990-2017. Watersheds were considered used if one or more wild pig location(s) were present. Available watersheds were randomly selected at a 1:1 ratio with used watersheds within the study area of the convex hull of wild pig occupied watersheds, with no overlap occurring between used and available watersheds. Model fit was validated using k-fold cross selection with a subset of the dataset that was withheld (10%) (Boyce et al., 2002). Biotic and abiotic covariates included in the model were identified from the literature across both the native and introduced ranges of wild pigs. However, as little is known regarding wild pig habitat and resource use at the northern extent of their introduced range, an exploratory approach to developing *a priori* models was used.

Cover type covariates were calculated as the proportion of each cover type within each watershed. The minimum distance to point locations of wild boar farms, large and small-scale domestic pig farms, and paved roads were calculated in meters in ArcGIS 10.6 (ESRI Redlands CA., 2018). Landscape heterogeneity within watersheds was measured using Simpson’s Diversity Index (McGarigal, 2015) through the program FRAGSTATS (University of Massachusetts, 2002). Variables were screened for variance inflation factors (VIF) and collinearity using Pearson’s correlation. Variable(s) with the least explanatory power were removed if two or more variables were highly correlated (rs>0.6 or VIF>0.5) (Zurr et al., 2007). All variables were standardized on a scale between 0 and 1 by subtracting the mean and dividing by the standard deviation to allow for comparisons between different scales and models (Zurr et al., 2007).

*A priori* models were developed from the set of biotic and abiotic variables and generalized linear models were used to analyze the data in R 3.5 (R Core Team, 2018) using the ‘MuMIN’ package (Barton, 2019). An information theoretic approach to model selection was used, in which inferences from a set of *a priori* models are based on a set of averaged models rather than a single best model (Burnham and Anderson, 2004). Multi-model inference is based on a set of models, that when within Δ2AIC are averaged to allow for inferences to be made based on the set of models that best describe the relationship of the data (Blakenship et al., 2002; Burnham and Anderson, 2002). The information theoretic approach is beneficial for use with exploratory analyses, as it is rare that a single top model best describes the patterns occurring in the data, especially when models in the set are closely related (Burnham and Anderson, 2002). The benefit of multi-model inference increases when a particular variable is present in all models (Burnham and Anderson, 2002). Variable importance in multi-model inference can be derived from Akaike weights (wi), which provide each variable in the set of models with an estimate of relative variable importance by summing the weight for a given variable across all models in the set where it occurs (Burnham and Anderson, 2002; 2004). As a result, all variables in a model set can be ranked by their individual relative importance *w+(j*) (Burnham and Anderson, 2002). The objective of the RSF was to quantify and characterize wild pig resource selection across the Prairie Provinces and to identify the relative risk of spatial overlap between wild pigs and domestic pig farms. As such, instead of using watersheds solely within the wild pig convex hull study area to visualize the RSF output, all domestic pig farm occupied watersheds across the Prairie Provinces were used to display the RSF output, which was slightly outside the wild pig study area. The relative risk of wild pig presence within a domestic pig farm occupied watershed was estimated based on the available habitat and resources present within each watershed. An RSF output was created for each classification of domestic pig farm watersheds: all farms combined, large-scale operations, and small-scale operations.

### 2.5 Invasive Wild Pig Visitations to Domestic Pig Farms

Wild pigs were captured by net-gun fired from a helicopter (n = 33) and using corral-style traps (n = 5) from 2015 to 2017 in southeastern and north-central Saskatchewan. Each wild pig was physically restrained and fitted with a Global Positioning System (GPS) tracking collar (Telonics, Mesa Arizona, USA). Collars were programmed to record a location every three hours and transmit the data via Iridium satellite link. GPS collars were intended to remain viable for one year, however, collar failures, losses, and hunter kills occurred throughout the one-year time period. As such only collars that collected a full year of data were retained and used in this analysis (n=11). The location accuracy of the collars was 6m and the fix rate success was 90% (Kramer, 2021). Wild pig data collection was approved by local ethical review [SPECIFIC DETAILS REMOVED FOR DOUBLE BLIND PEER REVIEW].

The GPS collar data were analyzed in two ways. Initially, the data was pooled to examine the overall trends in the distance to domestic pig farms by wild pigs. Data was pooled by the geographic location where the collars were deployed; 6 individual wild pigs for the Moose Mountain (MM) area in southeastern SK (females n=4, males n=2) and 5 individual wild pigs for the St. Brieux (STB) area in the north-central area of SK (females n=2, males n=3). The collar data from the STB study area was collected over a one-year time period from 2016-2017. Five of the collars from the MM study area collected data over a one-year time period from 2016-2017. One collar collected data over a one-year time period from 2015-2016. The one-year time period was separated into six two-month seasons to generate accurate proportions of seasonal data. The distance (km) from each GPS point to the nearest farm was calculated in ArcGIS 10.6 (ESRI Redlands CA., 2018) and was evaluated in R 3.5 (R Core Team, 2018) as a factor of season (early winter, late winter, spring, early summer, late summer, and autumn), month, time (AM/PM), sex, and farm type (large, small). T-tests and one-way ANOVA tests were used to determine if there were any significant differences (p<0.05) in the distance to domestic pig farms per factor.

Secondly, the fine-scale movements of individual collared wild pigs in the MM study area were analysed to determine if any discernible patterns or trends were detected in wild pig visitations to domestic pig farms. The distance (km) of 6 individual wild pigs (females n=4, males n=2) to domestic pig farms was measured and examined as a factor of; i) season; ii) time of day; and iii) individual domestic pig farms (Farm ID) in southeastern Saskatchewan. Five of the collars collected data over a one-year time period from 2016-2017, while one collar collected data over a one-year time period from 2015-2016. One-way ANOVA tests were calculated to determine if there were any significant seasonal or diurnal differences (p<0.05) observed in the distance of wild pigs to domestic pig farms and whether the distance to individual domestic pig farms differed.

Visits by wild pigs to domestic pig farms were quantified as a direct measure of spatial overlap between domestic and wild pigs. Farm visitations were classified as a UTM location recorded from a GPS collar within a 1-km buffer around a domestic pig farm. The three-hour interval fix rate of the GPS collars (Kramer, 2021) leaves gaps in wild pig movement data, therefore the 1-km buffer accounts for wild pig movement within the buffered areas in close proximity to farms that may have been missed (Wycoff et al., 2009). Furthermore, the foraging and wallowing behaviour of wild pigs may contaminate soil and water (Payne et al., 2016), increasing the chance of indirect disease transmission on non-biosecure operations via fomites and contaminated resources. Hence, a larger buffer captures a greater amount of risk in terms of indirect contact potential and provides results that can be interpreted for all types of domestic pig farms. The proportion of seasonal visitations of the 6 collared wild pigs to large-scale domestic pig farms (n=8) over a one-year period were evaluated in the Moose Mountain study area. Chisquare goodness of fit tests were calculated to determine if there were significant differences (p<0.005) in the proportion of wild pig seasonal visitations (≤ 1km) to Farm 1448. Only one domestic pig farm, Farm 1448, was considered in this analysis, as the number of visitations by wild pigs to the other farms in the study area were too low to conduct a chi-square test.

### 2.6 Risk Assessment of Wild Pigs and Domestic Pig Farms

A risk assessment was developed to estimate the likelihood of disease transmission from wild to domestic pigs using spatial overlap as a proxy for disease transmission. Multiple different datasets were used (wild pig distance (km) to domestic pig farms, the distance (km) between domestic wild boar and domestic pig farms, farm type, and landcover variables: proportion annual crop, proportion deciduous forest, and proportion water) in the analysis to characterize the spatial and environmental parameters and domestic pig farm demographics within the study area. Incorporating multiple datasets enabled the creation of a relative scale of risk to domestic pig farms for the potential of disease transmission from wild pigs. Separate risk assessments were completed for all farms combined, large-scale domestic pig farms, and small-scale pig farms.

All datasets were converted to raster data and continuous and integer data was standardized to a comparable scale between 1 and 10. Using the weighted sum tool in ArcGIS 10.6 (ESRI, Redlands, CA., 2018), each raster dataset was weighted according to its influence on the presence of wild pigs from 1 (low) to 3 (high). Parameter weights were allocated through statistical inference from the RSF models, as well as knowledge obtained via field experience and through the literature. The weighted sum tool multiplies the value of the given weight with the value of each raster cell for each parameter and sums all rasters together to create an overall scale of relative risk.

The parameters distance to wild boar farms and distance of wild pigs to domestic pig farms were accorded the highest weights in the model. The distance to wild boar farms had one of the largest parameter estimates in the RSFs determined by Aschim and Brook (2019). The distance of wild pigs to domestic pig farms provides a logical concept of risk, as wild pigs closer to farms provide an increased measure of potential risk to domestic pigs. Distance was measured between point locations instead of at the watershed level to provide a finer spatial scale measure of distance. Wild pig and wild boar farm locations across all years (1990-2017) were used, as wild pigs do not disperse far from the initial sources of introduction (wild boar farms) onto the landscape. Distance was measured in meters in ArcGIS 10.6 (ESRI, Redlands, CA., 2018) and converted to kilometers.

Three landcover variable raster datasets were used in the weighted overlay analysis - annual crop, deciduous forest, and water. These three variables were statistically significant in the RSF from Aschim (2022) as they comprised the largest beta values and are considered universally important drivers of wild pig presence throughout the literature (Beasley et al., 2014; McClure et al., 2015; Froese et al., 2017; Lewis et al., 2017; Snow et al., 2017). Landcover raster data were weighted so as to having moderate importance in the overlay analysis. Landscape heterogeneity also had high beta values in the RSF, however, optimal wild pig habitat contains a mixture of annual crop, deciduous cover, and water. Therefore, the use of the three most important variables indirectly represents landscape heterogeneity and accounts for its importance in wild pig presence on the landscape. The variables were mapped as the proportion of each variable type per watershed unit.

The variables farm type and farm density were weighted with the lowest importance in the weighted overlay. Farm density consisted of the number of farms per watershed for each separate analysis (large scale, small scale, and all farms). Each class was reclassified by dividing the highest density of farms by 10 to standardize farm density across all three classes. Farm type was classified as either large or small scale and were arbitrarily reclassified with small-scale domestic pig farms having a higher classification (i.e., more risk). The likelihood of direct and indirect contact between domestic and wild pigs is greater for small-scale farms as opposed to large-scale domestic pig farms due to the lack of biosecurity measures.

## 3. Results

### 3.1 Wild Pig and Domestic Pig Farm Distribution Mapping

A total of 2,549 domestic pig farms, within 894 watersheds (area = 331,542.7 km^2^) were present across the study area (Table 1). The total area of large and small-scale domestic pig farms was 204,282 km^2^ and 220,284 km^2^ respectively, however the density within watersheds was 8.5 times greater for large-scale domestic pig farms. Spatial overlap of wild pigs with all, large-scale domestic pig farms, small-scale domestic pig farms, and domestic wild boar farms was 20.7%, 20.9%, 20.6%, and 52.9% with the average distance from farm centroids to the nearest wild pig point locations at 42.2 km (SD 1.44), 42.3 km (SD 2.12), 38.2 km (SD 1.58), and 16.52 km (SD 19.10) respectively.

**Table 1.** Summary of the area (based on the Level 9 watershed unit) and number of domestic pig farms across Alberta, Saskatchewan, and Manitoba, 2019. Wild boar farms include point locations of all historical and current farms across the study area from 1990-2017. The overlap of domestic pig farms with wild pig occupied watersheds and the minimum, maximum, and mean distance of domestic pig farms, calculated as the centroid of the legal land description location, to wild pig point locations was calculated for all, large, and small-scale domestic pig farms, as well as domestic wild boar farms across the study area.

| Farm Type | Number of Domestic Pig Farms | Total Area (km<sup>2</sup> Watersheds) | Area (km<sup>2</sup>) Overlap Farms with Wild Pig Watersheds | % Overlap Farms with Wild Pig Watersheds | Mean Distance (km) Between Farm Centroid and Wild Pig Location | Min Distance (km) Between Farm Centroid and Wild Pig Location | Max Distance (km) Between Farm Centroid and Wild Pig Location |
| --- | --- | --- | --- | --- | --- | --- | --- |
| All Domestic Pig Farms | 2,549 | 337,084.33 | 69,812.18 | 20.71 | 42.22 | 1.04 | 255.28 |
| Large-Scale Pig Farms | 1,649 | 204,282.10 | 42,597.34 | 20.85 | 47.27 | 1.04 | 255.28 |
| Small-Scale Pig Farms | 900 | 220,284.71 | 45,319.44 | 20.57 | 38.22 | 1.04 | 222.15 |
| Domestic Wild Boar Farms | 127 | 45,308.25 | 23,516.27 | 52.94 | 16.52 | 0.00 | 84.93 |
| Wild Pig Occupied Watersheds |  | 131,709.26 |  |  |  |  |  |

Used wild pig locations were significantly closer to domestic pig farms (t=34.74, p<0.001, mean distance used points = 12.6km) and domestic wild boar farms (t=45.8, p<0.001, mean distance used points = 25.7km) as compared to random points on the landscape. Wild pigs were significantly closer to small-scale (mean distance = 10.6 km) domestic pig farms than large-scale (mean distance = 17.7 km) domestic pig farms (t= 8.8, p <0.001). The distance of wild pigs to wild boar farms (mean distance=6.7 km) was significantly closer than the distance of wild pigs to domestic pig farms (mean distance = 10.3 km, t= −6.4, p<0.001). Results of the non-linear regression showed that the number of wild pig occurrences was greatest within 0-20 km of domestic pig farms and decreased linearly as distance increased (R2 = 0.94, p<0.001).

### 3.2 Pathogen Presence and Wild Pig Overlap

The Canadian distribution of wild pigs had considerable spatial overlap with areas of *M. bovis* (6,002.2 km^2^) and CWD (156,159.1 km^2^) established in wildlife populations (Fig. 2), with most of the overlap occurring in Saskatchewan (83%) and Alberta (15%). *M. bovis*, CWD, and Pseudorabies were established in wildlife populations in five U.S. states bordering Canada (Montana, North Dakota, Minnesota, Michigan, and New York) (Fig. 3). Wild pigs had established populations in provinces bordering infected states, or within the states themselves. PRRVS, *B. suis*, and Pseudorabies were present in wild pig populations in western, eastern, and southern U.S. states and had a widespread geographic distribution. African Swine Fever (ASF) has not been identified in Canada and so there was no overlap with invasive wild pigs.

**Figure 2.**
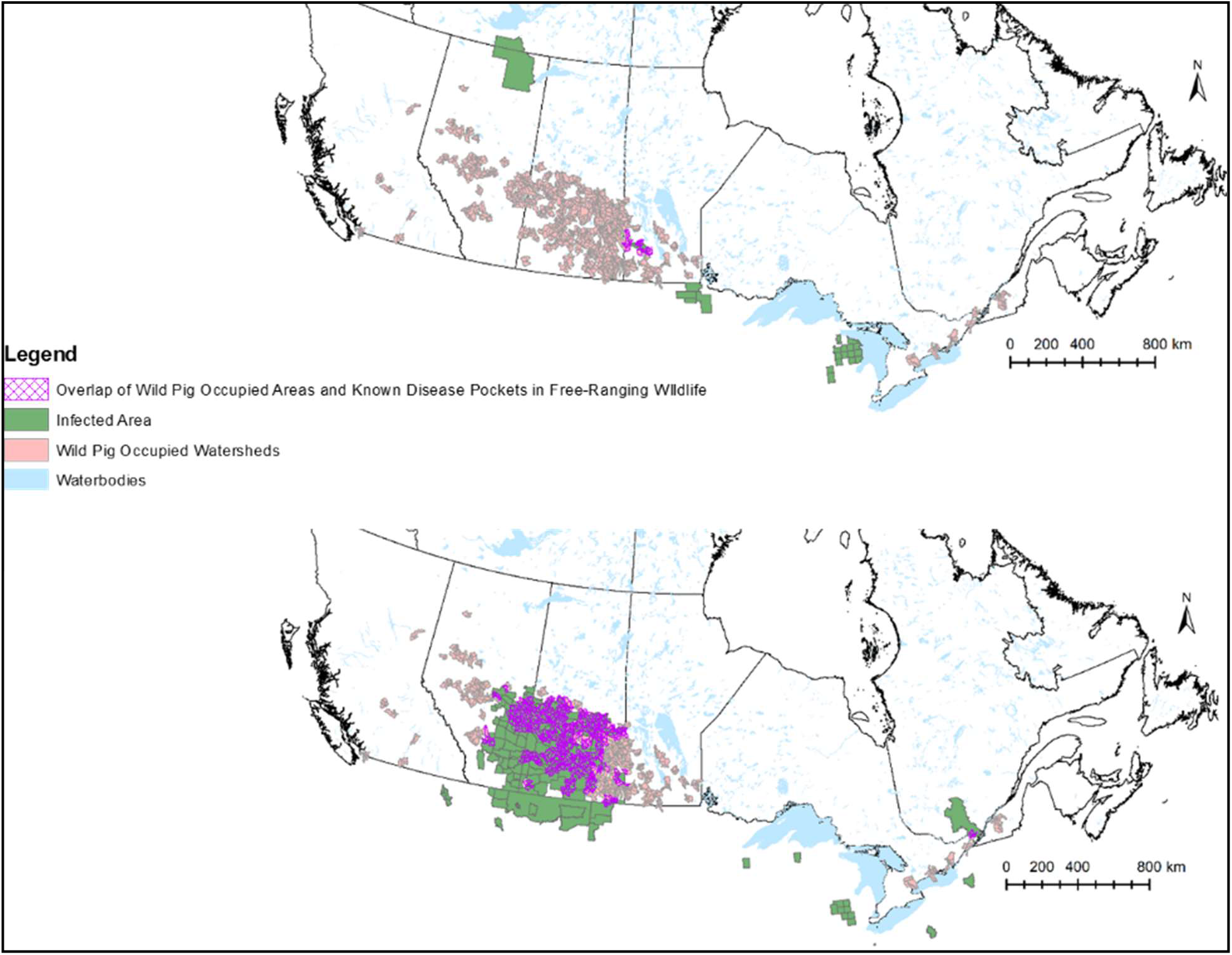
Spatial overlap of wild pigs, mapped at the Level 9 watershed unit, and areas of a)*Mycobacterium bovis*, mapped at the National Park (Canada) and county (U.S.) level, and b) CWD, mapped at the wildlife management zone unit (Canada) and the county (U.S.) level, established in wildlife populations in Canada and northern U.S. states, 2017-2019.

**Figure 3.**
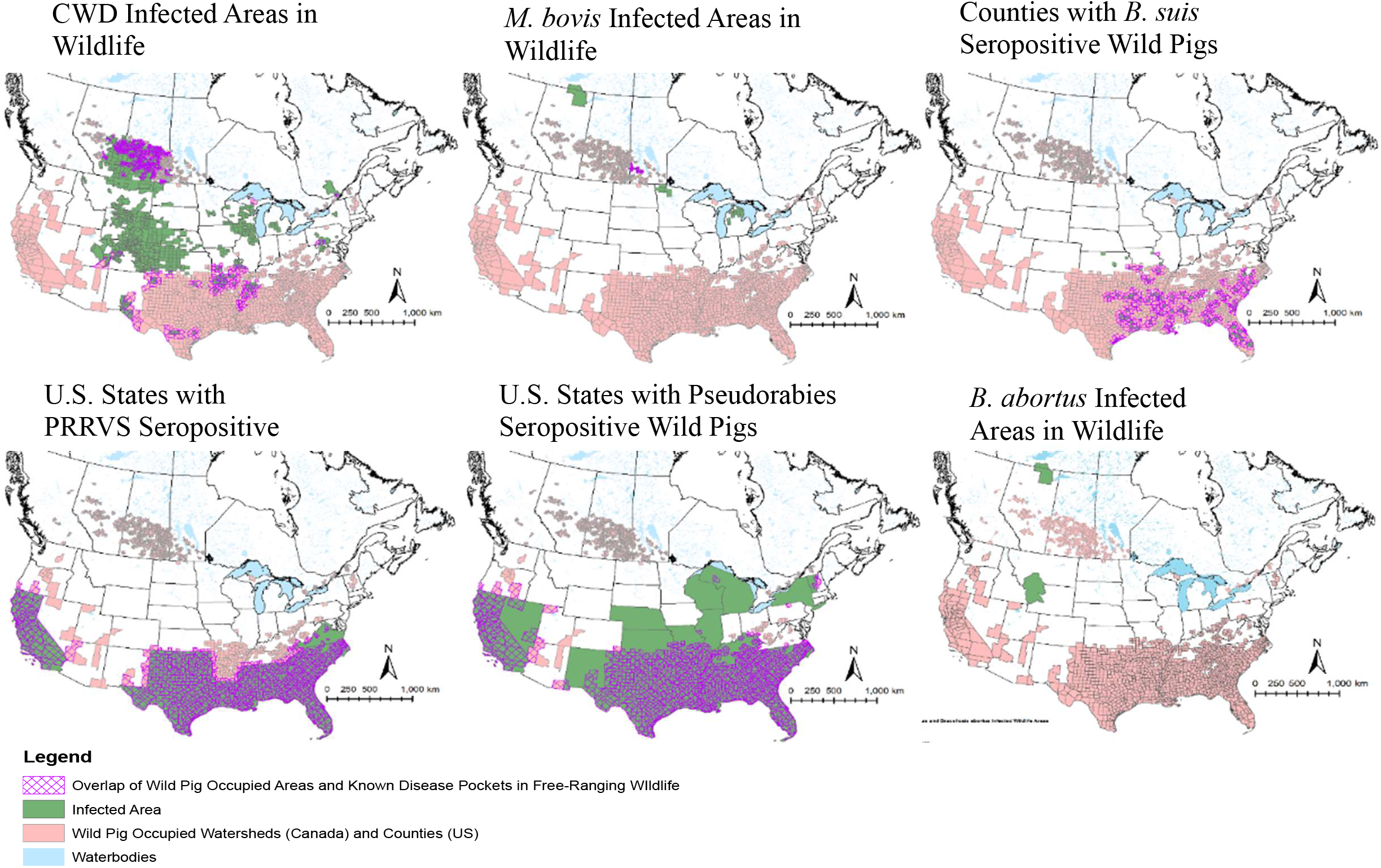
Spatial overlap of wild pigs in Canada, mapped at the Level 9 watershed unit, and the U.S., mapped at the county level unit, with areas of CFIA reportable diseases established in wildlife and wild pig populations, 2017-2019. Pathogen data was obtained from provincial and state governments. U.S. wild pig locations were obtained from state governments.

### 3.3 Habitat and Resource Use of Invasive Wild Pigs

The distance to wild boar farms had the greatest influence on wild pig distribution across the landscape from the model averaged results of the RSF; (β= −1.22 [−1.54, −0.93]; Fig. 4). Additional variables positively associated with wild pig presence on the landscape included annual crop cover (β [95% CI] = 0.614 [0.21, 1.19]), deciduous forest cover (β [95% CI] = 0.449 [−0.05, 1.03]), water (β [95% CI] = 0.107 [−0.29, 1.0]), landscape heterogeneity (β [95% CI] = 0.511 [−0.001, 0.87]), and distance to roads (β [95% CI] = −0.324 [−0.68, −0.01]) (Tables 2 and 3). Binomial logistic regression models of individual variables showed that wild pigs displayed a linear response to habitat and resource selection. Use increased as the proportion of annual crop cover increased and gradually decreased as the proportion of deciduous forest cover and water increased (Fig. 5). Wild pigs also displayed a positive linear response to perennial cover; however, the variable perennial was removed from the model(s) due to correlation with annual crop cover. Wild pigs avoided coniferous forest cover (β [95% CI] = −0.912 [−1.56, 0.18]), wetlands (β [95% CI] = −0.006 [−0.49, 0.23], and developed areas (β [95% CI] = −0.391 [−1.06, 0.50]). Use by wild pigs decreased linearly as the proportion of coniferous forest cover and wetlands increased.

**Figure 4.**
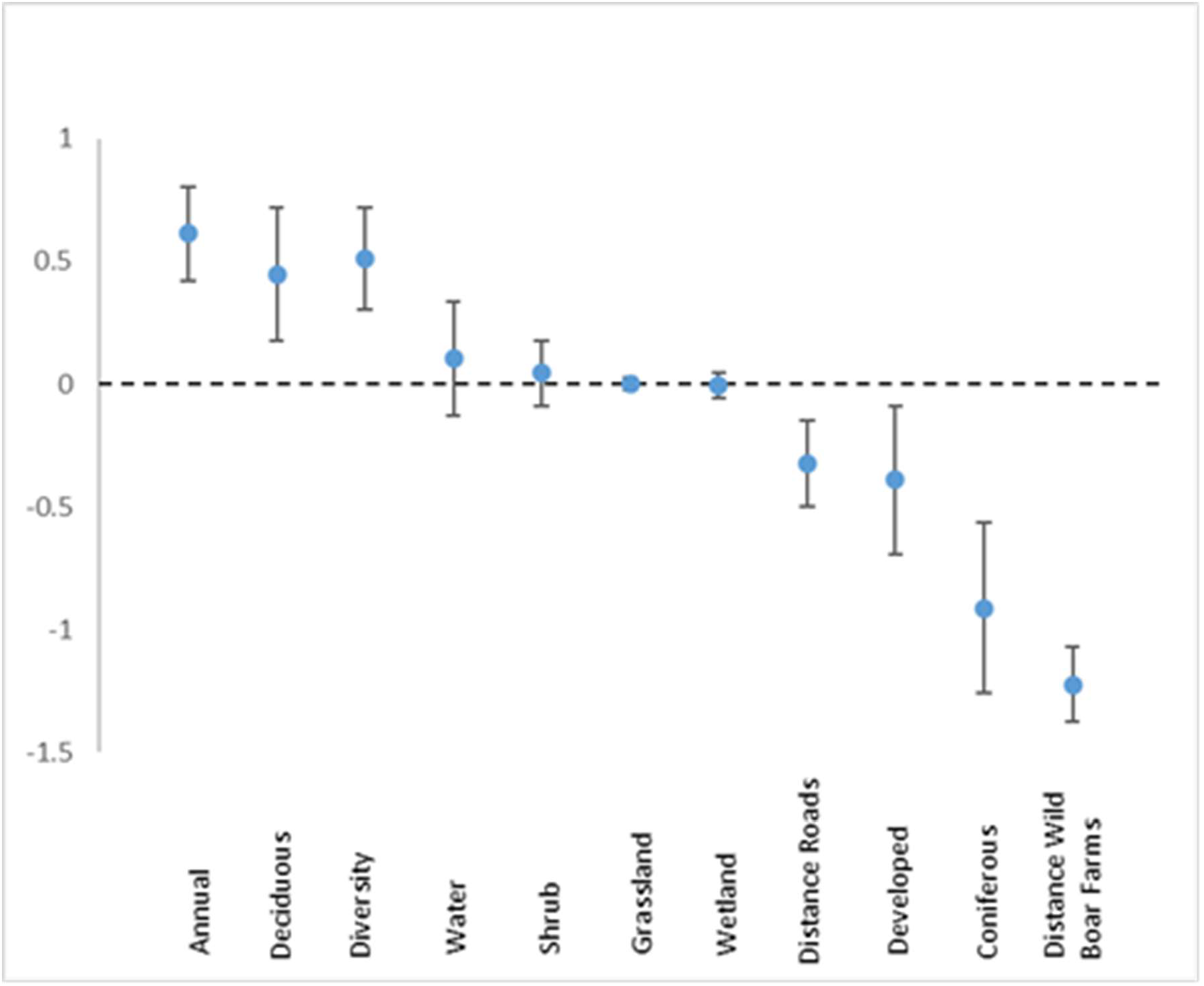
Model averaged beta coefficients and standard errors of biotic and abiotic predictor variables from the top models of the wild pig resource selection function analysis at the Level 9 watershed scale 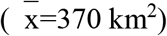 in the Prairie Provinces, 1990-2017.

**Figure 5.**
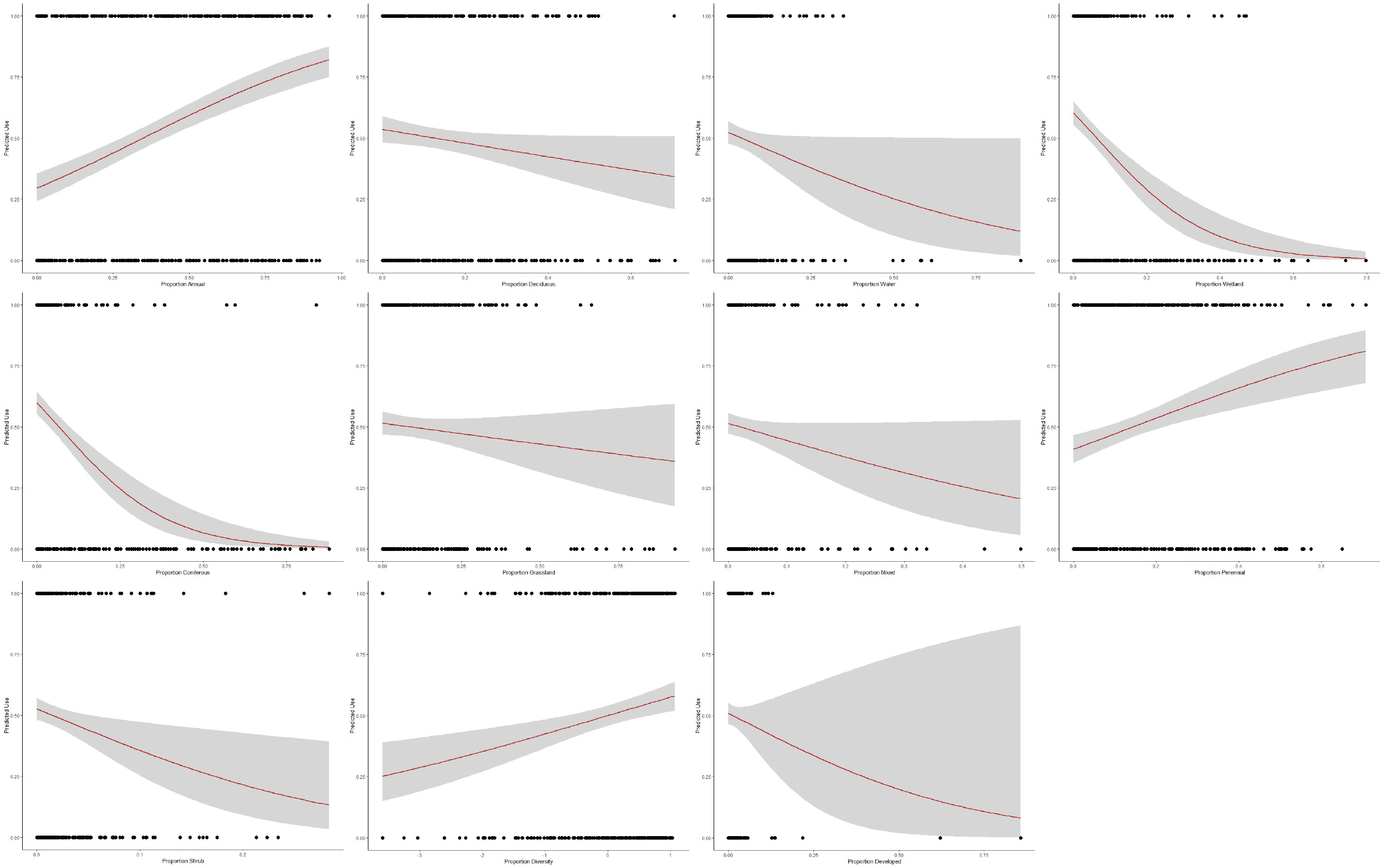
Plots of binomial logistic regression models for each cover type variable used in the RSF model. The plots depict the predicted use of each cover type by wild pigs between 0 and 1 (0 = low likelihood of use, 1 = high likelihood of use) and 95% confidence intervals based on the cover type proportion within watersheds 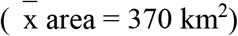 across the Prairie Provinces.

**Table 2.** Model averaged beta coefficients, standard errors, and Akaike weights [w+j] for predictor variables from the top models of the wild pig resource selection function analysis within domestic pig farm occupied watersheds across the Prairie Provinces 1990-2017.

| Variable | $\beta$ | S.E. | [w+j] |
| --- | --- | --- | --- |
| Annual | 0.614 | 0.190 | 1.00 |
| Deciduous | 0.449 | 0.274 | 0.91 |
| Coniferous | -0.912 | 0.348 | 1.00 |
| Water | 0.107 | 0.234 | 0.25 |
| Diversity | 0.511 | 0.208 | 1.00 |
| Shrub | 0.044 | 0.133 | 0.42 |
| Wetland | -0.006 | 0.049 | 0.06 |
| Grassland | 0.002 | 0.027 | 0.03 |
| Developed | -0.391 | 0.301 | 1.00 |
| Dist Boar Farms | -1.22 | 0.153 | 1.00 |
| Dist Roads | -0.325 | 0.176 | 0.97 |

**Table 3.**
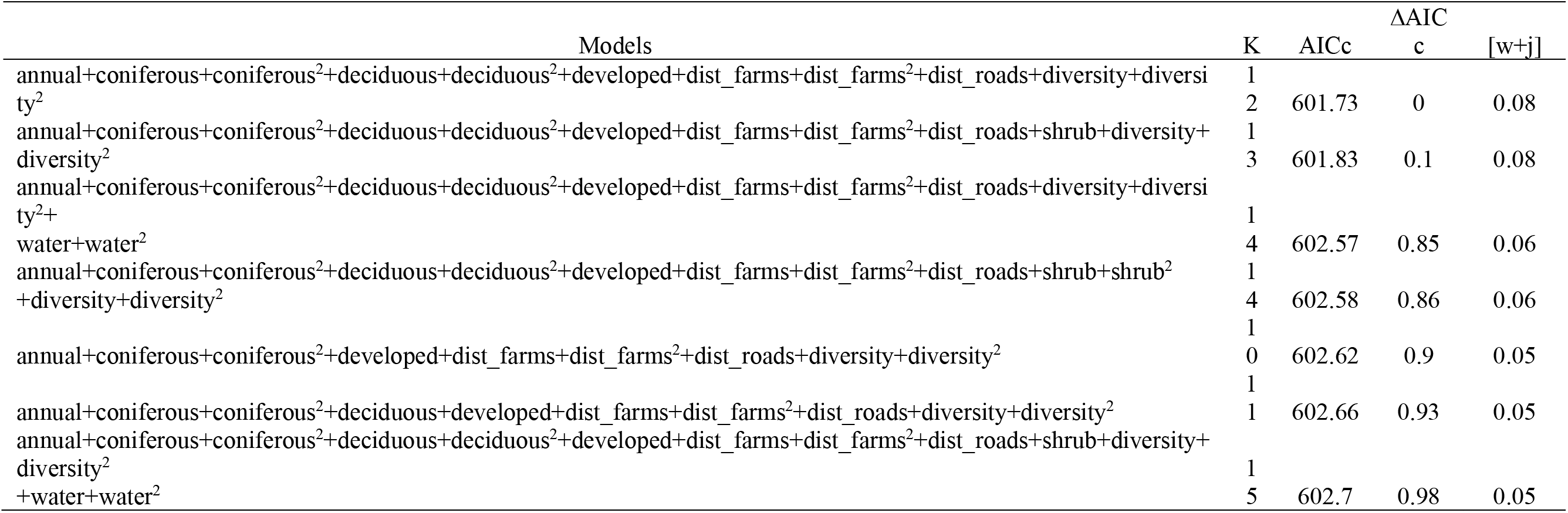
Number of model parameters, Akaike’s Information Criterion for small sample sizes (AICc), difference in AICc values from the top model (ΔAICc) and Akaike weights [w+j] for the top seven models of the wild pig resource selection within domestic pig farm occupied watersheds across the Prairie Provinces, 1990-2017.

### 3.4 Invasive Wild Pig Visitations to Domestic Pig Farms

The distance to domestic pig farms by wild pigs was significant for the factors sex (MM t=- 52.33, df=15091, p<0.001; STB t=-48.91, df=12827, p<0.001), farm type (MM t=-106.82, df=9295, p<0.001; STB t=93.70, df=1785, p<0.001), month (MM F=71.03, df=11, p<0.001; STB F=473.9, df=11, p<0.001), and season (MM F=55.71, df=5, p<0.001; STB F=888.7, df=5, p<0.001) for the pooled data in the MM and STB regions. Distance to domestic pig farms was not significant for the factor time of day for either study area (MM t= −0.14, df=18521, p=0.87; STB t=1.89, df=12839, p=0.06). The difference in the results between the two geographic locations, however, was such that no trends in wild pig distance to domestic pig farms were observed between the two geographic locations.

The six collared wild pigs within southeastern SK were within a distance of 0-16 km of large-scale domestic pig farms throughout the one-year time period. The average distance of all 6 pigs to domestic pig farms was 5.3 km (females 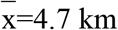, males 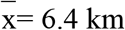). Individual wild pigs displayed significant differences in the average distance to domestic pig farms between seasons, however no overall trends in seasonal visitations to domestic pig farms were observed between individual wild pigs. Significant differences in the average distance of wild pigs to individual farms were observed for all individual pigs. No significant differences were observed in the average daily distance (day vs night) of wild pigs to domestic pig farms. Significant differences (p<0.001) in the seasonal visitations to Farm 1448 were observed for each individual pig. However, the proportion of seasonal visitations to Farm 1448 and all farms within the MM study area varied between pigs and no overall trend(s) were observed. Within the 1km buffer, the mean distance of wild pigs to a domestic pig farm was 435m (Fig. 6).

**Figure 6.**
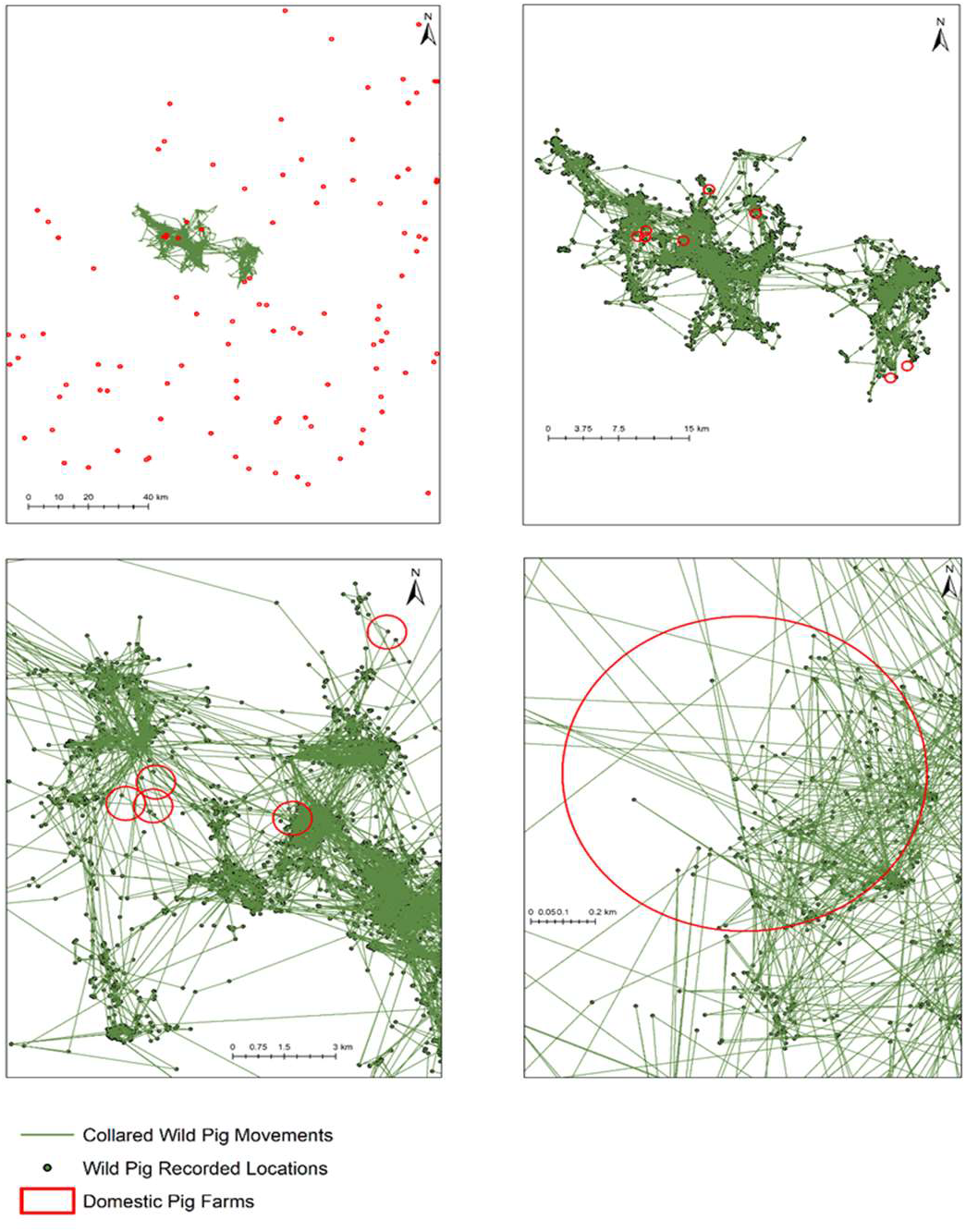
Visualization of wild pig visitations to domestic pig farms. Farms are buffered by 1km and the pooled three-hour interval GPS locations from wild pig collars are connected to illustrate wild pig movement in southern Saskatchewan. Any wild pig location ≤1km is considered a farm visitation with spatial overlap occurring between wild pigs and domestic pig farms. The four panels illustrate the a) locations of domestic pig farms in the area surrounding collared wild pigs; plot b) scales down to the immediate area of the GPS collar locations. Eight domestic pig farms are within the geographic area of the wild pig GPS locations. Plot c) focuses on four domestic pig farms that had the majority of wild pig visitations: and plot d) zooms into the farm with the greatest number of wild pig visitations within one year (2016-2017). Location details have all been removed to maintain confidentiality.

### 3.5 Risk Assessment of Invasive Wild Pigs and Domestic Pig Farms

The weighted sum of cover type proportions, wild pig distance to domestic pig and wild boar farms, farm type, and farm density identified the relative risk of wild pig presence associated to each domestic pig farm occupied watershed (Fig. 7). The range of risk values in the risk assessment model were similar for all three risk assessments. The relative risk values were greatest for small-scale domestic pig farms (low= 21.61, high = 106.06) and lowest for large-scale domestic pig farms (low = 17.4, high = 102.81). The scale of relative risk of all domestic pig farms combined ranged from 17.33 to 105.71.

**Figure 7.**
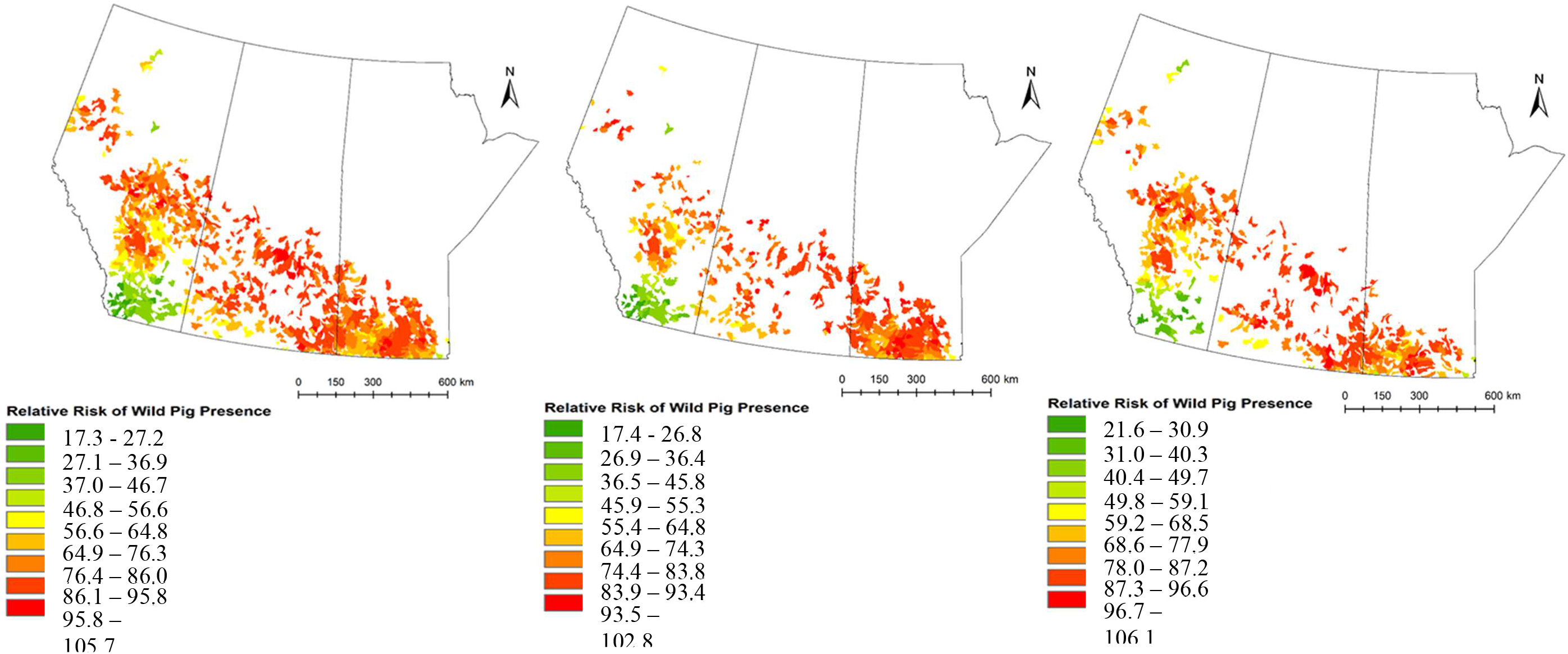
Relative scale of risk to domestic pig farms at the Level 9 watershed scale based on the biotic and abiotic covariates included in the weighted sum analysis. Relative risk is measured for domestic pig occupied watersheds for a) all domestic pig farms, b) large-scale domestic pig farms, and c) small-scale domestic pig farms in Alberta, Saskatchewan, and Manitoba. High values indicate a high risk of wild pig presence within domestic pig farm occupied watershed.

## 4. Discussion

The relative risk of spatial overlap between wild and domestic pigs is high across the Canadian Prairies. The potential for spatial overlap is driven by the landscape composition within domestic pig farm occupied watersheds and the distribution of current and historical domestic wild boar farms. High-quality wild pig habitat is located throughout domestic pig farm occupied watersheds across the study area, indicating the high relative likelihood of wild pig presence and future spread into these areas. The RSF model quantified the relative risk of spatial overlap between wild and domestic pigs as a measure of high-quality wild pig habitat available within domestic pig farm occupied watersheds. The use of predictive modeling has been widely used as a proxy measure to estimate the potential risk of disease transmission at the wildlife-livestock interface (Brook and McLachlan, 2009; Proffitt et al., 2011; Pruvot et al., 2014).

The results of the wild pig RSF and risk assessment are consistent with other studies where environmental variables identifying the relative probability of selection are identified as the most important proxy measure in predicting the relative likelihood of spatial overlap between wildlife and livestock over other factors such as farm characteristics and demographics and wildlife density (European Food Safety Authority, 2015; Carrasco-Garcia et al., 2016; Bosch et al., 2017).

Wild pig presence on the landscape is driven by landscape heterogeneity where an optimal mix of annual crop, deciduous forest, and water are available. This mix of habitat provides access to energy-rich agricultural food resources in proximity to cover and muddy wallows near water, which is consistent with wild pig habitat selection in other areas of their introduced ranges (Gaines et al., 2005; Paolini et al., 2018; Lewis et al., 2019). Wild pigs strongly avoided coniferous forest cover; however, this may be a factor of the distribution of wild boar farms across the study area in areas with low or no coniferous forest cover present and the minimal presence of coniferous forest within domestic pig farm occupied watersheds. We hypothesized that wild pigs would display a positive response to wetlands, however a negative response was observed. The negative response observed could partially be a result of the sparse wetland coverage in the spatial data layers, as many ephemeral or seasonal wetlands may not be defined. There may also be a trade-off observed in the landscape-level analysis that was applied for the RSF model. Whereas the use of a large-scale model is more conducive to management and planning (Hobbs, 2003; Johnson et al., 2005), fine-scale cover types, such as individual wetlands, may either be lost at the larger grain sizes or may not be selected for by the individual based on the individual’s perceptual range (Laforge et al., 2015) at the larger extent.

The strong positive response to distance from wild boar farms in the model indicates that wild pigs are not dispersing far from initial introduction points. As in Aschim (2022), the significance of the parameter estimate of distance to wild boar farms in the landscape scale model indicates that the current distribution of wild pigs across the Prairies is largely driven by introduction events. The spatial distribution of current and historical wild boar farms, and as a result introduction events, are in areas of high agricultural land use and domestic livestock production. Therefore, the geographic location of initial introduction events and slow outward spread from these points facilitates spatial overlap with domestic swine. The variables distance to domestic pig farms; all farms, large-scale, and small-scale farms were included in the *a priori* models, however the variables were insignificant, therefore were not included in the final models. The insignificant results from the domestic pig farm variables indicated that domestic pig farms were not influencing wild pig distribution and movement across the landscape. Rather, the initial sources of introduction coupled with available habitat and resources were the driving factors of wild pig distribution across the Prairies.

Although considerable overlap of wild pigs and infected wildlife is occurring in Canada, we are not aware of any testing that has been conducted to identify the presence or prevalence of any of the CFIA reportable or annually notifiable diseases in wild pigs. The limited presence of CFIA reportable diseases established in Canadian wildlife populations suggests that the risk of pathogen transmission into wild pig populations and subsequently to domestic pigs is likely generally low. However, this remains still largely unclear as testing of wildlife, especially wild pigs remains very limited. Overlap between wild pigs and *M. bovis* infected cervids in the Riding Mountain National Park (RMNP) area presents an opportunity for disease transmission of *M. bovis* into wild pig populations, though recent testing has shown a dramatic decrease if not eradication of bTB in wild cervids in and around RMNP (Canadian Food Inspection Agency, 2019). However, bovine tuberculosis is notorious for being difficult to completely eradicate, particularly in areas with diverse wildlife and livestock hosts.

The spatial overlap of wild pigs and Chronic Wasting Disease (CWD) infected cervids across the Prairie Provinces was widespread. In addition to the spatial overlap identified in the analysis, CWD has recently been identified in Manitoba (Government of Manitoba, 2021), further demonstrating the difficulties in managing the spread of disease in wildlife populations. Although spatial overlap between wild pigs and CWD infected areas has been identified, a relatively high species barrier for CWD transmission exists between cervids and suids (Moore et al., 2017). However, a recent study by Moore et al., (2017) found that pigs orally and intracranially inoculated with CWD brain homogenate from infected cervids had detectable disease associated prion proteins (PrP^sc^) in the brain and lymphoid tissues. Therefore, the potential for wild pigs to become infected with CWD is present, and transmission potential exists through rooting and feeding of contaminated vegetation and soil (Moore et al., 2017). Furthermore, since wild pigs are omnivores and are active scavengers, they are likely to feed on carcasses of CWD infected cervids (Wilcox and van Vuren, 2009; McDonough et al., 2022). Wild pigs may be able to disperse the prions across the landscape even if the species themselves do not become infected. As such, wild pigs have the potential to act as CWD reservoirs that are capable of shedding the disease for long periods of time before they develop clinical symptoms (Moore et al., 2017), which poses new risks for increased CWD spread.

In the southern U.S. wild pigs are hosts and reservoirs of PRRVS (Pedersen et al., 2018), *B. suis* (Olsen, 2010; Pedersen et al., 2012), and Pseudorabies (Corn et al., 2004; Pedersen et al., 2013). Although a large geographic distance currently exists between infected populations of wild pigs in the U.S. and the Canadian wild pig population, the potential for disease introduction to Canada exists through illegal translocations and the northward range expansion of wild pigs in the U.S. Infected wild pigs and other wildlife species in U.S. states bordering Canada pose serious risks for disease spread into Canada. A large concern is the presence of Pseudorabies infected wild pigs in U.S. states bordering Canada. Pseudorabies is not currently present in Canada; however wild pigs are present in provinces bordering Pseudorabies infected states which creates increased potential for disease introduction into Canada. African Swine Fever (ASF) is a swine disease that is highly contagious and can cause catastrophic economic impacts to domestic pig production. ASF has never been detected in Canada or the United States, but it is found in many countries around the world and is widespread in domestic and wild pigs in Asia and Europe. It has been detected in North America in the Dominican Republic and Haiti. It is single most important current disease of concern to the domestic swine sector in Canada and is relevant since it is well established that ASF can be spread between wild and domestic pigs (Sauter-Louis et al. 2021).

Our relative risk assessment identified small-scale domestic pig farms as having the greatest risk of spatial overlap with wild pigs, thus, the highest potential risk of disease introduction, spill-over, and spill-back events. Overall, 35% of domestic pig farms across the study area are small-scale backyard or hobby farms with variable biosecurity measures. These operations are widespread across the study area and direct overlap of small-scale domestic pig farms and wild pig occupied watersheds occurs throughout the study area. Although biosecurity measures vary on a per-farm basis, the lack of biosecurity measures more commonly found on small-scale operations allows for direct contact between individuals as well as indirect contact through access to feed and water. Within our study area we are also aware of at least two cases of mating between a free-ranging wild pig and a domestic pig on farms that resulted in hybrid offspring (Aschim pers. obs.). Disease transmission from direct and indirect contact between wild pigs and domestic swine on farms with no biosecurity measures has been identified in the U.S. and Europe (Wyckoff et al., 2009; Bosch et al., 2017; Anderson et al., 2019). Within the study area, 65% of domestic pig farms were large-scale biosecure operations. The risk of disease transmission from wild to domestic pigs at these facilities is relatively low due to the high biosecurity measures in place. However, risk is not zero as some diseases can be transmitted indirectly through aerosol transmission via droplets through ventilation systems (Corn et al., 2009). If strict biosecurity measures are not adhered to the potential of transmission from fomites, such as contaminated clothing, footwear, equipment, and infected feed is present (Lowe et al., 2014; Pasick et al., 2014; Kim et al., 2017; Alarcon et al., 2021). Importantly, large domestic pig barns can be strong attractants to wild pigs both through spilled feed and odours from barns, especially for boars, where sows in heat may attract wild pigs from large distances (Wycoff et al., 2009; Brook et al., 2013; Pruvot et al., 2014).

The current map of wild pig spatial overlap with domestic pig farms identified within this study should be viewed as conservative estimates, as we recognize it was not possible to detect all wild pig occurrences and wild pigs in the Canadian Prairie Provinces continue to expand out of control so these maps are already in need of updating (Aschim and Brook, 2019). Additionally, the available data does not capture all small-scale domestic pig operations across the study area, as sporadic inventory of operations by the industry occurs and small-scale hobby farms may not be reported or are missed (Mark Ferguson, pers. comm.; Jeff Clark, pers. comm.). As such, the current spatial overlap we have mapped for wild and domestic pigs is now much greater than identified in this study.

The distance to domestic pig farms by wild pigs was significant for the factors sex, farm type, month, and season for the pooled data in the MM and STB regions. Within both study areas, females were within a closer distance to domestic pig farms than males, which was unexpected. This result may be somewhat biased due to the small sample size of collared individuals as well as farm type biases. Similar results were found in a study in Texas, therefore the greater visitations to farms by females could in fact be a behavioural bias due to the social behaviour of wild pigs (Wycoff et al., 2009). While the significant differences in the results are interesting, larger sample sizes in terms of individuals as well as sexes and a stratified sampling approach in terms of domestic pig farm type is required. Wild pigs in Saskatchewan have been found to have some of the largest home ranges within both their introduced and native ranges. The summer home ranges of wild pigs in Saskatchewan range from 100.98 km^2^ and 6.61 km^2^ at the 99% and 50% Brownian Bridge utilization distribution respectively (Brook et al., unpublished). Therefore, although no overall trends were observed in terms of the seasonal visitations of wild pigs to domestic pig farms, the large home ranges observed within the study area signifies that at multiple different spatial scales, the average distance of wild pigs to domestic pig farms is well within the seasonal movements of wild pigs. The lack of observed trends identifies the need for year-round mitigation measures.

All farms within the vicinity of GPS-collared wild pigs were large-scale biosecure operations. Therefore, farm visitations were likely a factor of the available habitat and resources rather than a preference for farm type/co-mingling with domestic pigs. The insignificance of the domestic pig farm variables in the models and the lack of any trends in the visitations to domestic pig farms by wild pigs signified that the driving factor in wild and domestic pig spatial overlap was the landscape composition and the location of domestic wild boar farms. However, this study was limited by sample size and study location as the data analysis for farm visitations was developed post hoc of the sampling design and implementation. GPS collars were initially deployed to monitor wild pig home range and seasonal movements on the landscape. As such, the study design and location were not chosen with respect to farm characteristics or distribution, hence the seasonal and diurnal movements of wild pigs in the vicinity of small-scale domestic pig farms were not captured. The GPS collars were operational for one year, however collar failures and collared pig mortality occurred, and as such a limited number of collars collected data over a one-year period. Inferences on wild pig farm visitation characteristics as a result cannot be definitively made but suggest several important areas for future study. Additional research is required regarding the diurnal and seasonal visitations of wild pigs to small-scale operations, and the characteristics of visitations. The recent optimization of GPS collars with video cameras built in would contribute critically to better understanding how wild pigs interact with domestic pigs and habitats on and near domestic pig farms.

As a relatively new invasive species on the landscape, wild pigs are continuing to expand their range across the Prairie Provinces. The large amount of high-quality wild pig habitat and resources available across the Canadian Prairies indicates the capacity for future spread into new areas. Continued and uncontrolled spread of the species further increases the current and potential risks of spatial overlap between wild and domestic pigs. A high relative likelihood of spatial overlap is estimated across the majority of the study area. Furthermore, overlap is expected to increase due to the widespread and expanding distribution of wild pigs in an agriculture dominated landscape, the amount of high-quality wild pig habitat within the study area, and the high number of domestic pig farms at moderate densities within watersheds occupying a broad geographic area. Similar trends have been observed in the U.S. in recent years (Wormington et al., 2019). Indeed, the study area is situated in the Prairie Provinces where wild pig populations are well-established and where the majority of the species range expansion in Canada (92%) has occurred (Aschim and Brook, 2019). Therefore, continued and increased spatial overlap and co-mingling between wild and domestic pigs is expected.

The current overlap of wild and domestic pigs across the study area and the average distance of wild pigs to domestic pig farms provides producers and the industry with the ability to implement cost-effective mitigation efforts. Invasive wild pigs are now firmly established with the study area and complete eradication of wild pigs in Canada is no longer possible, creating significant long-term challenges. The year-round spatial overlap observed coupled with the insignificant results of the diurnal visitations signifies that specific seasonal and diurnal mitigation measures for spatial overlap between wild and domestic pigs are ineffective and year-round mitigation is required. Camera traps on farms are an effective means of obtaining large amounts of data at a low cost which can provide further insight into the seasonal and diurnal visitations to farms by wild pigs. Enhanced on-farm mitigation measures include robust fencing, wildlife deterrents, and population control within a buffer distance around farms (Wycoff et al., 2009). Further research and continued monitoring are required on the spatial movements and farm visitation characteristics of wild pigs, particularly in regards to small-scale domestic pig farms.

The results highlight the role wild pigs would play in driving the spread of disease and the obstacle they present in disease eradication in livestock. Diseases at the livestock-wildlife interface present complex elimination and control challenges, therefore risk analysis and mitigation strategies that prevent wildlife-livestock interaction and/or overlap are central to the prevention of wildlife-livestock disease transmission (Miller et al., 2013). Routes of transmission between wildlife and livestock and the ability of wild pigs to act as natural reservoirs of disease are still poorly understood (Witmer et al., 2003; Miller et al., 2013; Miller and Sweeny, 2013; De la Torre et al., 2015). Consequently, all routes of transmission should be identified and taken into consideration when designing on-farm mitigation and prevention measures.

Disease testing of wild pigs has been conducted in a small part of the study area (SK), and individuals have been relatively healthy (McGregor et al., 2015). However, testing of the range of CFIA reportable and annually notifiable diseases was not included and testing of wild pigs was opportunistic with no rigorous sampling in place. Furthermore, the known overlap of wild pig populations with *M. bovis* and CWD infected wildlife provides an increased and ever-present risk of disease transmission to the swine industry. Increased and targeted surveillance and monitoring of pathogen presence in wild pig populations is required, specifically in proximity to domestic pig farms, as early detection is an extremely effective management tool that can be utilized as a preventative measure (Miller et al., 2013).

Wild pig habitat and resource use was modeled at a broad scale to coincide with the scale at which management decisions and planning are made (Hobbs, 2003). Domestic pig farms and wild pigs are widely distributed across the Canadian prairies, therefore regional-scale analyses, surveillance, mitigation, and disease control planning is required. The generalist nature of the species lends itself well to analyses at large spatial scales. Broad habitat and land-use types are appropriate for making inferences and can adequately address the mechanisms influencing land use by wild pigs, hence the subsequent risks to farms. The broader scale of analysis also provides results that allow for direct application towards future management and risk planning to address the complex social-ecological dimensions associated with wild pigs (Hobbs, 2003; Froese et al., 2017).

Domestic pig farms are widespread across the study area and differ in terms of farm characteristics, demographics, and biosecurity measures. Therefore, specific management and mitigation measures need to be implemented on a per-farm basis and a detailed on-farm risk assessment should be conducted to complement our large-scale analysis. The results of this risk assessment analysis identify priority areas for increased disease surveillance, enhanced on-farm mitigation measures, and focused efforts toward wild pig population control. The findings can be directly applied to management planning to allow for quick detection and rapid response should a disease outbreak occur.

